# Canonical Wnt-signaling modulates the tempo of dendritic growth of adult-born hippocampal neurons

**DOI:** 10.1101/2020.01.14.905919

**Authors:** Jana Heppt, Marie-Theres Wittmann, Jingzhong Zhang, Daniela Vogt-Weisenhorn, Nilima Prakash, Wolfgang Wurst, Makoto Mark Taketo, D. Chichung Lie

## Abstract

In adult hippocampal neurogenesis neural stem/progenitor cells generate new dentate granule neurons that contribute to hippocampal plasticity. The establishment of a morphologically defined dendritic arbor is central to the functional integration of adult-born neurons. Here, we investigated the role of canonical Wnt/β-catenin-signaling in dendritogenesis of adult-born neurons. We show that canonical Wnt-signaling follows a biphasic pattern, with high activity in stem/progenitor cells, attenuation in early immature neurons, and re-activation during maturation, and demonstrate that the biphasic activity pattern is required for proper dendrite development. Increasing β-catenin-signaling in maturing neurons of young adult mice transiently accelerated dendritic growth, but eventually resulted in dendritic defects and excessive spine numbers. In middle-aged mice, in which protracted dendrite and spine development was paralleled by lower canonical Wnt-signaling activity, enhancement of β-catenin-signaling restored dendritic growth and spine formation to levels observed in young adult animals. Our data indicate that precise timing and strength of β-catenin-signaling is essential for the correct functional integration of adult-born neurons and suggest Wnt/β-catenin-signaling as a pathway to ameliorate deficits in adult neurogenesis during aging.

## Introduction

In the adult mammalian hippocampus, neural stem/progenitor cells in the subgranular zone of the dentate gyrus undergo a complex sequence of proliferation, differentiation, and maturation steps to add dentate granule neurons to the hippocampal network. A key step for the functional integration of adult-born dentate granule neurons is the development of a morphologically highly stereotypic dendritic arbor to receive afferents in the molecular layer (Goncalves, Schafer et al., 2016b). Disruption of dendritic arbor development of adult-born neurons is thought to contribute to cognitive and emotional deficits in aging, neurodegenerative and neuropsychiatric diseases, and to the development of an aberrant circuitry in epilepsy (Cho, Lybrand et al., 2015, Fitzsimons, van Hooijdonk et al., 2013, Jessberger & Parent, 2015, Kerloch, Clavreul et al., 2018, Kim, Liu et al., 2012, Li, Bien-Ly et al., 2009, Llorens-Martin, Rabano et al., 2015, Murphy, Hofacer et al., 2012, Sun, Halabisky et al., 2009, Trinchero, Buttner et al., 2017, Winner, Melrose et al., 2011).

In young adult mice, the dendritic arbor of adult-born dentate granule neurons is largely established within the first 3-4 weeks of development. Around 10 days after their birth, new neurons feature a basic dentate granule neuron architecture with an apical dendrite spanning the dentate granule cell layer and initial branching in the inner molecular layer. Dendritic growth with further branching and dendritic extension into the outer molecular layer is maximal during the first two to three weeks and is followed by a period of pruning of excessive dendritic branches to attain the highly stereotypic dentate granule neuron morphology (Goncalves, Bloyd et al., 2016a, Kleine Borgmann, Bracko et al., 2013, Sun, Sailor et al., 2013, Zhao, Teng et al., 2006). Several factors including hippocampal network activity, transcription factors, cytoskeletal regulators, neurotransmitters, and signaling molecules were found to modulate dendrite morphology of adult-born neurons (Bergami, Rimondini et al., 2008, Gao, Ure et al., 2009, Ge, Goh et al., 2006, He, Zhang et al., 2014, Jagasia, Steib et al., 2009, Llorens-Martin, Fuster-Matanzo et al., 2013, Ma, Jang et al., 2009, Piatti, Davies-Sala et al., 2011, Trinchero et al., 2017, Vadodaria, Brakebusch et al., 2013). A complete understanding of the central pathways controlling dendrite development in adult hippocampal neurogenesis is, however, still missing.

Wnt proteins are key regulators of adult hippocampal neurogenesis (Arredondo, Guerrero et al., 2019, Jang, Bonaguidi et al., 2013, Lie, Colamarino et al., 2005, Qu, Sun et al., 2010, Qu, Sun et al., 2013, Seib, Corsini et al., 2013). Current data indicate that Wnts control different stages of adult neurogenesis via distinct pathways: while early developmental steps such as proliferation and fate determination of precursors are regulated by canonical Wnt/β-catenin-signaling (Karalay, Doberauer et al., 2011, Kuwabara, Hsieh et al., 2009, Lie et al., 2005, Qu et al., 2013), late developmental steps such as neuronal maturation and morphogenesis are thought to be primarily regulated by non-canonical Wnt-signaling pathways (Arredondo et al., 2019, Schafer, Han et al., 2015). Supporting a sequential action of distinct Wnt-pathways is the finding that early adult-born neuron development and the initiation of neuronal morphogenesis are accompanied by the attenuation of canonical Wnt/β-catenin-signaling activity and the increased activity of the non-canonical Wnt/planar cell polarity (PCP) signaling pathway (Schafer et al., 2015). However, the observations i) that pathologies associated with aberrant Wnt/β-catenin activity are paralleled by dendritic growth defects of adult-born dentate granule neurons (De Ferrari, Avila et al., 2014, Duan, Chang et al., 2007, Llorens-Martin et al., 2013, Martin, Stanley et al., 2018, Murphy et al., 2012, Qu, Su et al., 2017, Singh, De Rienzo et al., 2011), and ii) that ablation of β-catenin from developing dentate granule neurons in juvenile mice causes massive dendritic defects and neuronal death (Gao, Arlotta et al., 2007), raise the possibility that Wnt/β-catenin-signaling fulfills important functions during late steps of adult hippocampal neurogenesis.

We here report that attenuation of canonical Wnt-signaling in early immature neurons is followed by re-activation of the pathway during maturation, resulting in a biphasic pattern of canonical Wnt-signaling activity in the adult neurogenic lineage. We also show that this biphasic activity pattern is essential to ensure correct dendrite development and that canonical Wnt/β-catenin-signaling activity in maturing neurons modulates the tempo of dendritic growth and spine formation. Finally, we demonstrate that countering the age-associated decrease in canonical Wnt/β-catenin-signaling reverses dendritic growth and spine formation deficits of adult-born neurons in middle-aged mice. Thus, our data reveal a new cell-autonomous function of canonical Wnt-signaling in maturation of adult-born neurons and suggest canonical Wnt/β-catenin-signaling as a candidate pathway to counteract age-related deficits in hippocampal neurogenesis-dependent plasticity.

## Results

### Canonical Wnt-signaling exhibits biphasic activity during adult hippocampal neurogenesis

Analyses of different transgenic reporter mouse lines consistently revealed high activity of canonical Wnt-signaling in the adult dentate gyrus (DG) (Garbe & Ring, 2012, Lie et al., 2005, O’Brien, Harper et al., 2004) but were inconclusive regarding canonical Wnt-signaling activity during different stages of adult-born neuron development (Garbe & Ring, 2012). To shed light on the activity pattern of canonical Wnt-signaling in adult neurogenesis, we analyzed the adult neurogenic lineage in two different reporter mouse lines: BATGAL mice harbor the LacZ reporter gene downstream of seven TCF/LEF-binding sites and the minimal promoter-TATA box of the *siamois* gene (Maretto, Cordenonsi et al., 2003); whereas Axin2^LacZ/+^ mice heterozygously harbor the LacZ reporter in the endogenous locus of the bona fide canonical Wnt-signaling target Axin2 (Lustig, Jerchow et al., 2002) (Fig. 1A). Both lines showed a biphasic pattern of canonical Wnt-signaling activity (Fig. 1B and C). Moderate activity was observed in NESTIN+ radial glia-like stem/progenitor cells. Immature neurons that were identified by the expression of DCX rarely displayed reporter gene expression; whereas mature dentate granule neurons that were identified by the expression of Calbindin again showed high reporter gene expression. Reporter activity in BATGAL animals suggested moderate canonical Wnt-signaling activity in TBR2+ precursor cells, while reporter activity was readily attenuated in TBR2+ cells in Axin2^LacZ/+^ mice (Fig. 1B and C). These differences in the timing of attenuation of reporter activity likely arose from the different genetic setup of the reporter.

**Fig. 1.**
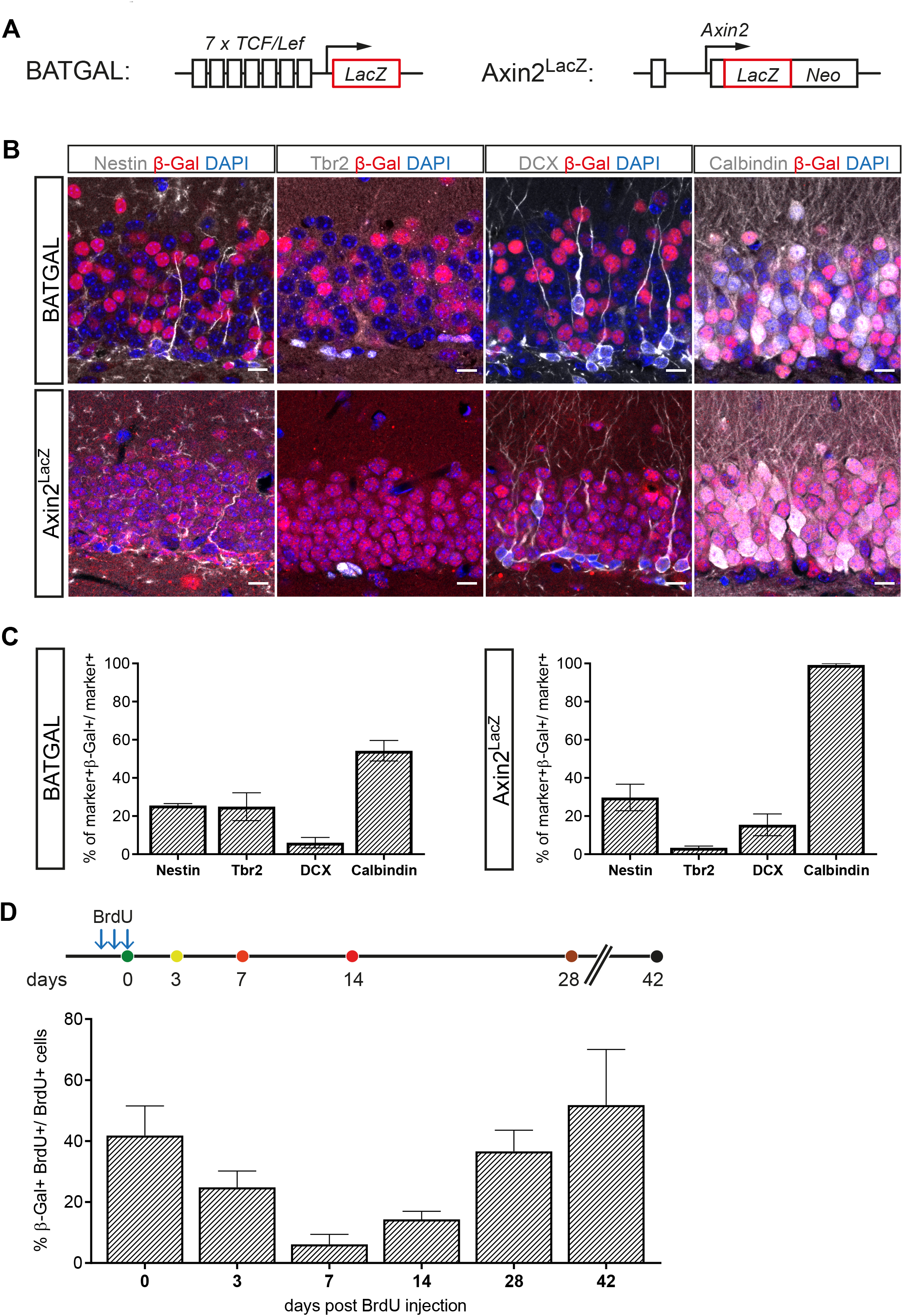
Canonical Wnt-signaling activity in the adult hippocampus. **A** Schematic representation of the *LacZ* alleles of the BATGAL and Axin2^LacZ^ mouse models for canonical Wnt-signaling activity **B** Representative images showing co-expression of the stage-specific markers Nestin, Tbr2, Doublecortin (DCX), and Calbindin, with the β-Galactosidase (β-Gal) reporter in 8-week old BATGAL and Axin2^LacZ^ mice. Scale bar=10μm **C** Fraction of Nestin-, Tbr2-, DCX- and Calbindin-positive cells expressing β-Gal in BATGAL and Axin2^LacZ^ animals showed stage specific canonical Wnt-signaling during adult hippocampal neurogenesis (n= 3 animals per mouse model and marker) **D** BrdU pulse chase scheme. BrdU was injected intraperitoneally (i.p.) three times: i) every two hours for 30-minute time point [0 days post injection (dpi)] ii) every 24 hours for all other time points. Animals were sacrificed 30-minutes after the final BrdU injection (0 dpi time-point; n= 4 animals), 3 dpi (n= 3 animals), 7 dpi (n= 3 animals), 14 dpi (n= 4 animals), 28 dpi (n= 6 animals) and 42 dpi (n= 6 animals). Quantification of cells co-expressing β-Gal and BrdU over all birthdated cells in the DG displayed biphasic activity of canonical Wnt-signaling during adult hippocampal neurogenesis. Data represented as mean ± SD.

To better assess the time course of canonical Wnt-signaling activity in adult neurogenesis, newborn cells in 8-week-old reporter mice were birthdated by a single pulse of Bromodeoxyuridine (BrdU). For this analysis we focused on the BATGAL reporter strain, because Axin2 functions in a feedback loop as a negative regulator of canonical Wnt-signaling (Lustig et al., 2002); consequently, heterozygous loss of Axin2 may affect the physiological developmental time course of adult-born neurons. BATGAL reporter activity in BrdU+ cells was analyzed at time-points that correspond approximately to the proliferating progenitor cell stage (thirty minutes post injection), the neuroblast stage [3 days post injection (dpi)], early and mid-immature neuron stage (7 and 14 dpi, respectively), and the early and late mature neuron stage (28 and 42 dpi, respectively) (Jagasia et al., 2009, Snyder, Choe et al., 2009). Thirty minutes after BrdU injection, 42% of BrdU+ cells showed reporter activity. This fraction dropped to 25% and 7% at the 3 dpi and 7 dpi time-points, respectively, to subsequently increase to 14% at 14 dpi, 38% at 28 dpi, and 52% at 42 dpi (Fig. 1D).

Collectively, these data show that canonical Wnt-signaling activity in the adult neurogenic lineage is attenuated in immature DCX+ neurons during the first week of development and is up-regulated around the second week during the maturation of DCX+ neurons into Calbindin+ neurons.

### Correct dendrite development of adult-born neurons is dependent on the timing and dosage of Wnt/β-catenin-signaling activity

We first asked whether the re-activation of canonical Wnt/β-catenin-signaling during maturation was required for adult-born neuron development. We focused in particular on dendrite development given that re-activation of canonical Wnt/β-catenin-signaling activity overlapped with the period of maximal dendritic growth speed (Sun et al., 2013). In canonical TCF/LEF transcription factor family (Moon, 2004). To inhibit canonical Wnt/β-catenin-signaling in the adult neurogenic lineage, we transduced fast-proliferating precursor cells with the CAG-dnLEF-IRES-GFP Mouse-Moloney Leukemia (MML) retrovirus, which bi-cistronically encodes for a dominant-negative LEF mutant protein (dnLEF) and GFP (Karalay et al., 2011). A MML retrovirus encoding for RFP (CAG-RFP) was co-injected with the CAG-dnLEF-IRES-GFP virus. Animals were analyzed 17 days post viral injection (Fig. 2A). Compared to RFP+ GFP-control neurons, CAG-dnLEF-IRES-GFP transduced neurons displayed a more immature morphology, with shorter total dendritic length likely caused by shorter terminal dendrites as indicated by the analysis of branch point number and Sholl analysis. In addition, dnLEF expressing neurons almost invariably exhibited basal dendrites, a transient feature of developing dentate granule neurons and morphological indication of immaturity (Ribak, Korn et al., 2004) (Fig. 2B - 2E). Hence, dnLEF-expressing neurons lagged behind control neurons with regard to dendrite development, indicating that canonical Wnt/β-catenin-signaling is required for the timely execution of the dendrite development program in adult-born dentate granule neurons.

**Fig. 2.**
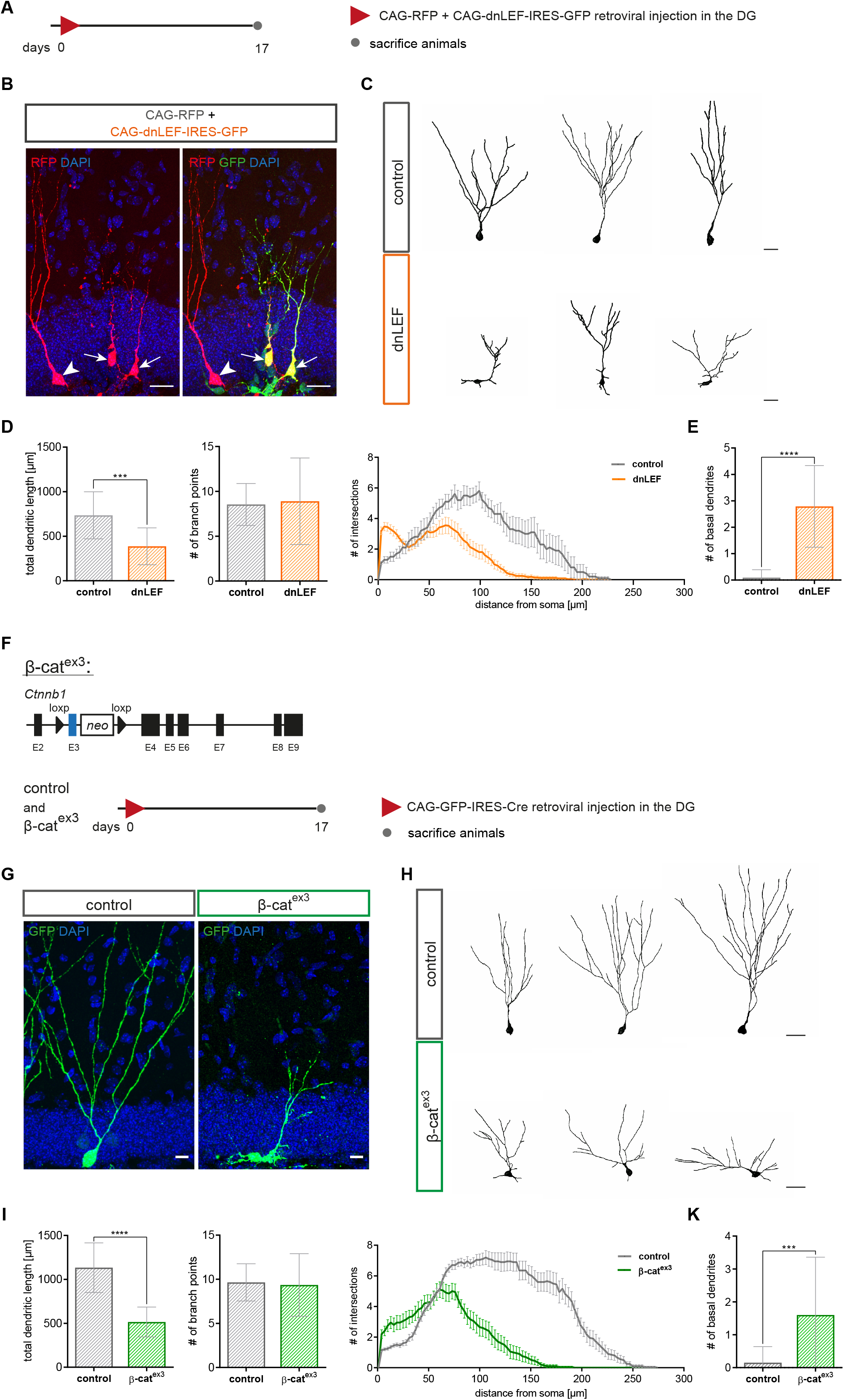
Loss of canonical Wnt-signaling and sustained canonical Wnt-signaling impair dendritogenesis of adult-born neurons. **A** Experimental scheme of retroviral injection paradigm. Adult mice were stereotactically co-injected with the MML retroviruses CAG-dnLEF-IRES-GFP (dnLEF) and CAG-RFP (control) and were sacrificed 17 days post injection (dpi). **B** Representative pictures of transduced adult-born neurons at 17 dpi. Arrows and arrowheads indicate CAG-dnLEF-IRES-GFP (dnLEF) and CAG-RFP (control) double transduced cells and CAG-RFP single transduced cells, respectively. Scale bar= 20 μm **C** Representative reconstructions of control and dnLEF neurons. Scale bar= 30 μm **D** Analysis of morphology showed a reduction in dendritic length in dnLEF transduced neurons, while number of branch points remained comparable. Sholl analysis displayed a reduction in dendritic complexity in dnLEF transduced neurons and indicated decreased growth of terminal dendritic branches [p<0.0001 (****)] (control: n= 11 cells from 3 animals, dnLEF= 38 cells from 6 animals). **E** DnLEF neurons also displayed basal dendrites (control: n= 11 cells from 3 animals, dnLEF= 38 cells from 6 animals). **F** Schematic representation of the conditional alleles of the Ctnnb1^(ex3)fl^ (β-cat^ex3^) mouse model and the retroviral paradigm used to analyze canonical Wnt-signaling gain of function in neural progenitors. Control animals harbor widtype allele for Ctnnb1. **G** Representative pictures depicting control and β-cat^ex3^ retrovirus transduced adult-born neurons at 17dpi. **H** Representative reconstructions of control and β-cat^ex3^ neurons. Scale bar= 30μm **I** Quantification showed decreased dendritic length, but no difference was apparent in number of branch points. Sholl analysis displayed a less complex dendritic tree (p<0.0001 (****))(control: n= 20 cells from 5 animals, β-cat^ex3^: n= 20 cells from 5 animals). **K** The number of basal dendrites was increased and the primary dendrite initiation site was slightly more horizontal (control: n= 20 cells from 5 animals, β-cat^ex3^: n= 20 cells from 5 animals). Data represented as mean ± SD (Sholl analysis: mean ± SEM), significance was determined using two-way ANOVA for Sholl analysis and two-tailed Mann-Whitney U Test for all other analyses, significance levels were displayed in GP style (p<0.0332 (*), p<0.0021 (**) and p<0.0002 (***), p<0.0001 (****)).

Given that inhibition of canonical Wnt/β-catenin-signaling retarded dendrite growth, we hypothesized that activation of canonical Wnt/β-catenin-signaling promotes dendritic development. The Ctnnb1^(ex3)fl^ mouse mutant (hereafter called β-cat^ex3^) allows for constitutive activation of canonical Wnt/β-catenin-signaling via Cre recombinase-induced expression of a stabilized form of β-catenin (Harada, Tamai et al., 1999) (Fig. 2F). To enhance canonical Wnt-signaling, we induced recombination in fast-dividing precursors by stereotactic injection of a MML retrovirus bi-cistronically encoding for GFP and Cre-recombinase (CAG-GFP-IRES-Cre). Ctnnb1^wildtype^ mice injected with the CAG-GFP-IRES-Cre retrovirus served as controls. Animals were analyzed 17 days post viral injection. Surprisingly, transduced neurons (GFP+; DCX+) in β-cat^ex3^ mice featured an immature dendritic arbor that was characterized by irregular short neurites, a decrease in total dendritic length and a reduced complexity in the Sholl-analysis (Fig. 2G – 2K), demonstrating that failure to attenuate Wnt/β-catenin-signaling by induction of activity of canonical Wnt-signaling in fast dividing neural progenitor cells disrupted physiological dendritogenesis.

Recombination of the β-cat^ex3^ locus and the resulting expression of stabilized β-catenin in fast dividing neural progenitor cells abolishes the early attenuation of canonical Wnt-signaling in the adult neurogenic lineage [Fig. 1 and (Schafer et al., 2015)]. To allow for an initial attenuation of canonical Wnt/β-catenin-signaling prior to genetic enhancement of pathway activity, we targeted recombination of the β-cat^ex3^ locus to a later developmental time-point, i.e., immature neurons. To this end, we generated a conditional β-catenin gain-of-function mouse line (DCX::CreER^T^² CAG-CAT-GFP; β-cat^ex3^ hereafter called β-cat^ex3^ iDCX) that allowed for tamoxifen-inducible expression of stabilized β-catenin in DCX-expressing immature neurons and for tracing of recombined cells by expression of a GFP reporter (Harada et al., 1999, Nakamura, Colbert et al., 2006, Zhang, Giesert et al., 2010). DCX::CreERT² CAG-CAT-GFP; Ctnnb1^(ex3)wt^ mice harboring wildtype alleles for Ctnnb1 served as controls. Recombination was induced in 8-week old, i.e, young adult mice by injection of tamoxifen on five consecutive days (Fig. 3A). Recombined neurons were identified by co-expression of GFP+ and the dentate granule neuron marker PROX1+. Analysis at 3 days after recombination showed that enhanced canonical Wnt/β-catenin-signaling activity stimulated the growth of terminal dendrite branches as evidenced by Sholl analysis, analysis of total dendritic length and number of branch points (Fig. 3B and C). We then asked whether enhanced dendrite growth would produce an excessive dendrite arbor. To this end, animals were analyzed 13 days after recombination. At this time point, β-cat^ex3^ iDCX neurons and control neurons showed a comparable branching pattern; surprisingly, β-cat^ex3^ iDCX neurons showed a trend towards decreased total dendritic length, while Sholl analysis revealed a reduction in the length of the terminal dendrites (Fig. 3D). These data suggest that increased Wnt/β-catenin-signaling only transiently accelerated dendrite growth and that enhancement of Wnt/β-catenin-signaling in adult-born neurons of young adult mice ultimately impeded dendrite development.

**Fig. 3.**
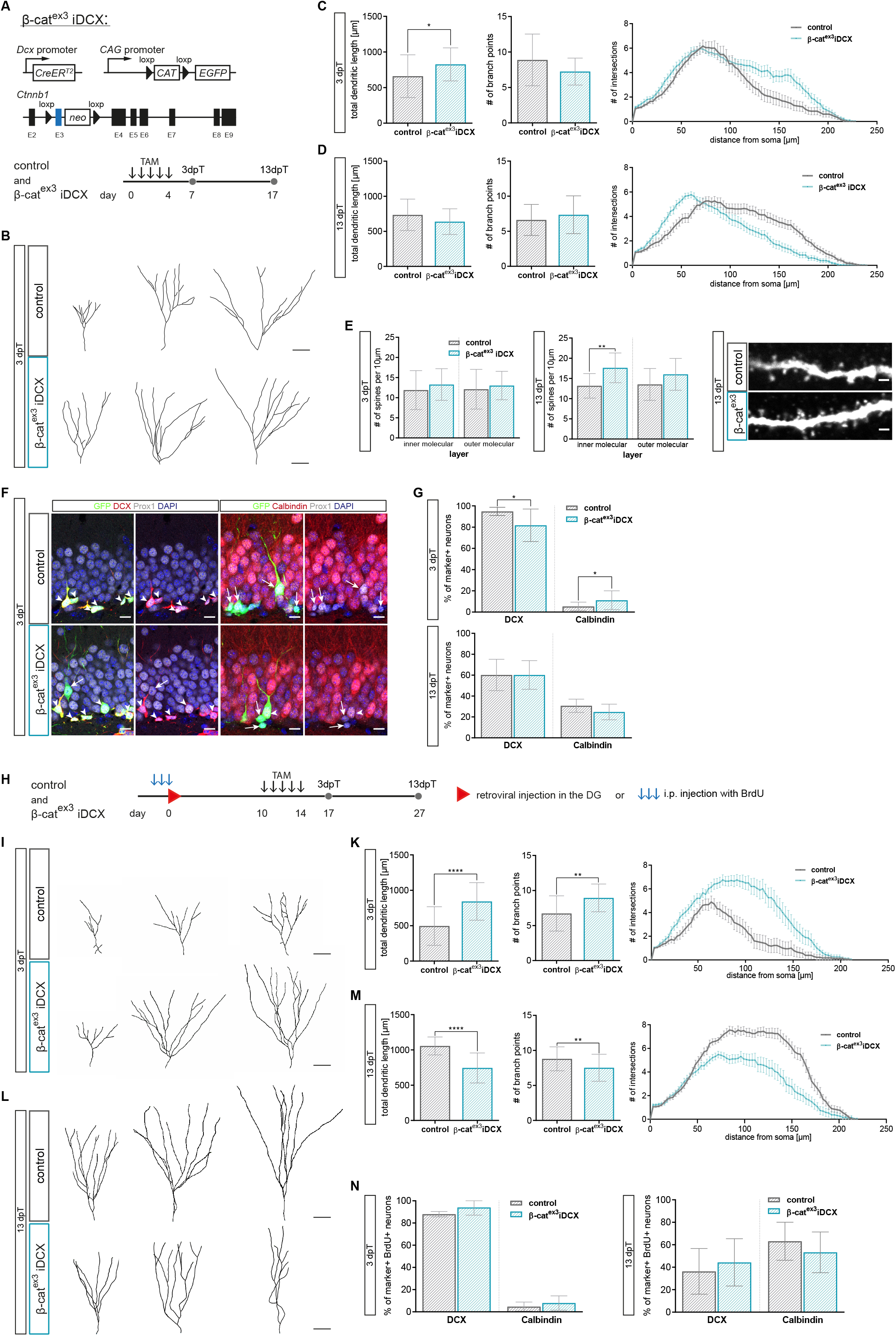
Enhanced canonical Wnt-signaling activity in immature neurons of young adult mice modulates dendritic growth and spine formation. **A** Schematic representation of the conditional alleles and transgenes of the β-cat^ex3^ iDCX mouse model. Tamoxifen (TAM) was applied i.p. every 12 hours for five days Animals were analyzed 3 and 13 days post Tamoxifen (dpT). **B** Representative reconstructions of control and β-cat^ex3^ iDCX neurons at 3dpT. Scale bar= 30μm **C** Analysis of morphology showed a slight increase of dendritic length in β-cat^ex3^ iDCX neurons at 3dpT. The number of branch points was unaltered. Sholl analysis indicated increased growth of terminal dendrite branches in neurons of β-cat^ex3^ iDCX mice [p<0.0001 (****) control: n= 25 cells from 9 animals; β-cat^ex3^ iDCX: n= 25 cells from 8 animals]. **D** Analysis of morphology showed no difference in dendritic length and branch point number between experimental groups at 13dpT. Sholl analysis indicated a decrease in length of terminal dendrites in neurons of β-cat^ex3^ iDCX mice [p<0.01 (**), control: n= 25 cells from 9 animals, β-cat^ex3^ iDCX: n= 25 cells from 8 animals]. **E** Spine densities were comparable between experimental groups at 3dpT. 13dpT β-cat^ex3^ iDCX neurons showed significantly increased spine density in the inner molecular layer (3dpT: control: n= 13 cells from 3 animals, β-cat^ex3^ iDCX: n= 13 cells from 4 animals; 13dpT control: n= 13 cells from 5 animals, β-cat^ex3^ iDCX: n= 16 cells from 7 animals). Representative images showing dendritic segments in the inner molecular layer of the DG at 13dpT. Scale bars= 1μm. **F** Representative pictures of recombined (GFP+) neurons co-expressing Prox1 as a marker for neuronal fate and the stage specific markers DCX for immature neurons and Calbindin for mature neurons. Arrows and arrowheads indicate marker negative and marker positive cells, respectively. Scale bar= 10μm **G** Fraction of DCX expressing neurons is slightly reduced in β-cat^ex3^ iDCX at 3dpT, while fraction of Calbindin expressing cells is slightly increased in β-cat^ex3^ iDCX at 3dpT. At 13dpT no difference is visible in stage specific marker expression (3dpT: control: n= 12 animals, β-cat^ex3^ iDCX: n= 12 animals, 13dpT: control: n= 10 animals, β-cat^ex3^ iDCX: n= 14 animals). **H** Experimental scheme for birthdating of adult-born neurons prior to recombination. For morphology analysis, CAG-RFP was stereotactically injected into the DG (I-M), for marker expression analysis, BrdU was injected i.p. every 24 hours for three days (N). Tamoxifen was applied i.p. every 12 hours for five days. Mice were sacrificed 3dpT and 13dpT. **I** Representative reconstructions of control and β-cat^ex3^ iDCX neurons expressing RFP at 3dpT. Scale bar= 30μm **K** Quantification of dendritic length and branch points, and Sholl analysis [p<0.0001 (****)] showed a higher dendritic complexity of β-cat^ex3^ iDCX neurons at 3dpT (control: n= 18 cells from 4 animals, β-cat^ex3^ iDCX: n= 19 cells from 5 animals). **L** Representative reconstructions of control and β-cat^ex3^ iDCX neurons expressing RFP at 13dpT. Scale bar= 30μm **M** Quantification of dendritic length, and branch points, and Sholl analysis [p<0.0001 (****)] showed a lower dendritic complexity of β-cat^ex3^ iDCX neurons at 13dpT (control: n= 20 cells from 4 animals, β-cat^ex3^ iDCX: n= 12 cells from 5 animals). **N** Expression of stage markers DCX and Calbindin in BrdU birthdated neurons was comparable between experimental groups at 3dpT and 13dpT (control: n= 5 animals, β-cat^ex3^ iDCX: n= 5 animals). Data represented as mean ± SD (Sholl analysis: mean ± SEM), significance was determined using two-way ANOVA for Sholl analysis and two-tailed Mann-Whitney U Test for all other analyses, significance levels were displayed in GP style [p<0.0332 (*), p<0.0021 (**) and p<0.0002 (***), p<0.0001 (****)]

To further evaluate the impact of enhanced β-catenin-signaling on neuronal maturation, we determined dendritic spine density as a proxy of the development of the glutamatergic postsynaptic compartment as well as the expression of the immature neuron marker DCX and the mature neuron marker Calbindin. While dendritic spine density was comparable between β-cat^ex3^ iDCX and control neurons at the 3 day time-point (Fig. 3E), β-cat^ex3^ iDCX neurons bore a higher dendritic spine density at the 13 day time-point (Fig. 3E) suggesting that canonical Wnt/β-catenin-signaling activity promoted spinogenesis. Three days after recombination 95% of recombined neurons (i.e., GFP+ Prox1+ cells) in control animals were DCX positive, while 5% expressed Calbindin. Activation of canonical Wnt-signaling resulted only in a small difference in the expression of maturation stage-associated markers: thus, β-cat^ex3^ iDCX animals showed a slightly decreased proportion of DCX+ immature neurons (82%) and a mildly increased fraction of neurons expressing Calbindin (11%) (Fig. 3F and G). Thirteen days after recombination, the fraction of DCX-expressing and Calbindin-expressing recombined neurons was comparable between control and β-cat^ex3^ iDCX mice (Fig. 3G). These data indicate that enhanced canonical Wnt/β-catenin-signaling activity in DCX+ neurons of young adult mice was not sufficient to substantially accelerate the maturation-associated switch from DCX-to Calbindin-expression.

The DCX::CreER^T2^ transgene targets canonical Wnt/β-catenin-signaling activity to immature neurons of various birthdates. To more precisely understand the impact of enhanced β-marker expression and dendrite parameters between neurons of a defined age. Neurons in β-cat^ex3^ iDCX and control mice were birthdated via BrdU-pulse labeling or transduction with a retrovirus encoding for RFP (CAG-RFP) prior to recombination. Animals received daily tamoxifen injections from day 10 until day 14 after birthdating. Consequently, BrdU-and RFP-labeled neurons were at most 14 days old at the time of recombination and thus were still in the phase of low or attenuated canonical Wnt-signaling activity. Animals were analyzed three and thirteen days after tamoxifen induced recombination (Fig. 3H). Analysis of BrdU-labeled recombined neurons (GFP+ BrdU+) indicated that increased β-catenin-activity did not affect the timing of the maturation associated switch from a DCX+/Calbindin-to a DCX-/Calbindin+ neuron (Fig. 3N). Three days after recombination, CAG-RFP birthdated neurons (GFP+ RFP+) with enhanced β-catenin activity bore a more mature dendritic arbor with substantially increased total dendrite length and a higher number of branch points compared to control neurons (Fig. 3I and K). As expected, control neurons showed substantial increase in dendritic length and complexity between the early and late time-point. In contrast, neurons in the enhanced β-catenin activity group did not display signs of further dendrite growth and refinement but rather showed a reduction in dendritic complexity to a level that was significantly below the level of control neurons (Fig. 3L and M). Thus, analysis of birthdated neurons confirmed that Wnt/β-catenin-signaling transiently accelerated dendrite growth but ultimately impeded dendrite development.

Collectively, our loss- and gain-of-function analyses demonstrate that physiological dendrite development of adult-born neurons is highly dependent on the timing and dosage of Wnt/β-catenin-signaling activity.

### Age-dependent decrease of canonical Wnt-signaling activity in the adult neurogenic lineage

Aging was reported to be accompanied by reduced Wnt production, activation of GSK3β and increased expression of Wnt-signaling inhibitors in the hippocampus (Bayod, Felice et al., 2015, Miranda, Braun et al., 2012, Okamoto, Inoue et al., 2011, Seib et al., 2013). To determine whether these age-associated alterations translate into decreased canonical Wnt-signaling in the dentate gyrus, we analyzed BATGAL mice at different ages. In young adult, 8-week old, BATGAL mice 49% of cells in the granule cell layer expressed the reporter β-Galactosidase. Previous studies have shown that adult neurogenesis in the murine dentate gyrus is already severely compromised around the age of 5 – 6 months (Ben Abdallah, Slomianka et al., 2010, Trinchero et al., 2017). Notably, the percentage of reporter positive cells was reduced to 26% in 24-week old mice, while in 36-week old mice only 21% of cells in the granule cell layer showed reporter activity (Fig. 4A).

**Fig. 4.**
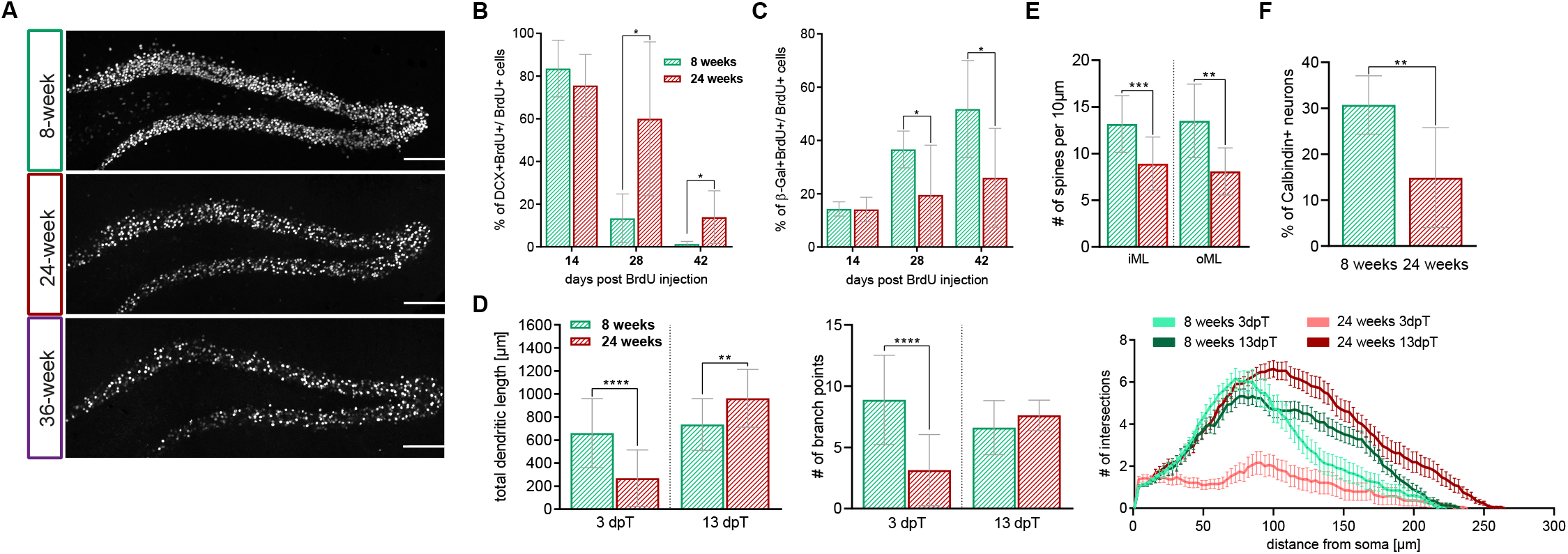
Canonical Wnt-signaling activity is decreasing with increasing age. **A** Representative overview images of the β-Galactosidase reporter signal in the dentate gyrus of 8-week old, 24-week old and 36-week old BATGAL mice. Note the decrease in reporter signal with increasing age. Scale bar= 100μm **B** Percentage of BrdU+ cells expressing DCX in young adult (8-week old) and middle-aged (24-week old) mice (14dpi: n= 4 animals, 18dpi: n= 6 animals, 42dpi: n= 6 animals). Note the prolonged expression of DCX in middle-aged mice. **C** Percentage of BrdU+ cells expressing the β-Galactosidase reporter in young adult (8-week old) and middle-aged (24-week old) mice (14dpi: n= 4 animals, 18dpi: n= 6 animals, 42dpi: n= 6 animals). Note the reduced reporter expression in middle-aged mice. **D** Comparison of total dendritic length, branch points, and Sholl analysis [p<0.0001 (****)] between 8-week-old and 24-week-old animals at 3dpT and 13dpT show age-associated delay in dendritic development (8 weeks: 3pT/13dpT n= 25 cells from 9 animals, 24 weeks: 3dpT/13dpT n= 21 cells from 5 animals). **E** Comparison of spine number in young and middle-aged mice showed reduced spine density in the inner and outer molecular layer of the dentate gyrus at 3dpT and 13dpT (8 weeks: n= 13 cells from 5 animals, 24 weeks: n= 14 cells from 4 animals). **F** Reduced percentage of neurons expressing the mature granule neuron marker Calbindin in middle-aged control mice compared to young adult control mice thirteen days after recombination (8 weeks: n= 10 animals, 14 weeks: n= 8 animals). Data represented as mean ± SD (Sholl analysis: mean ± SEM), significance was determined using two-way ANOVA for Sholl analysis and two-tailed Mann-Whitney U Test for all other analyses, significance levels were displayed in GP style [p<0.0332 (*), p<0.0021 (**) and p<0.0002 (***), p<0.0001 (****)].

Next, we investigated whether the age-associated decrease in canonical Wnt-activity also affected the neurogenic lineage. Adult-born cells in young adult (i.e., 8-week old) and middle-aged (i.e., 24-week old) BATGAL mice were birthdated with BrdU. In young adult mice, the fraction of BrdU+ cells expressing the immature marker DCX was high (82%) in 14-day old cells and dropped to 12% in 28-day old cells. 42 days after the BrdU pulse virtually all BrdU+ cells had lost DCX expression, implying that all cells had reached a mature phenotype 42 days after birth (Fig. 4B). Middle-aged mice featured a similar fraction of BrdU+ cells expressing DCX at the 14-day old time point; a higher percentage of cells, however, remained in the DCX+ stage at 28 days (58%) and 42 days (15%). The delayed exit from the DCX+ stage in middle-aged mice was paralleled by lower activity of canonical Wnt-signaling. 14 days post birthdating, the percentage of β-Galactosidase positive neurons among the BrdU-labeled cells was highly comparable between young and older mice mice (8-week old: 14%, 24-week old: 14%). While this percentage was increased in young adult mice at 28 dpi (37%), middle-aged mice exhibited only a negligible increase in β-Galactosidase expression (19%). At 42 dpi, 52% of BrdU-labeled cells in young mice expressed β-Galactosidase, while in older mice only 26% had active canonical Wnt-signaling (Fig. 4C). Collectively, these observations show that aging is associated with a substantial decrease of canonical Wnt-signaling in the adult neurogenic lineage.

### Canonical Wnt-signaling activity in immature neurons of middle-aged mice accelerates maturation

Aging protracts dendritic growth and spine development of adult-born neurons (Beckervordersandforth, Ebert et al., 2017, Trinchero et al., 2017). To determine whether increasing canonical Wnt-signaling could ameliorate the age-associated maturation defects of adult-born dentate granule neurons, we induced recombination in middle-aged (i.e., 24-week old) β-cat^ex3^ iDCX and control mice. Three days after recombination, neurons in 24-week old control mice displayed in comparison to neurons in 8-week old control mice a much simpler dendritic arbor with decreased dendritic length and lower number of branch points (Fig.4D). Thirteen days after recombination, age-associated differences in dendritic arbor morphology were no longer present (Fig. 4D). Neurons in older mice, however, showed substantially reduced spine densities (Fig. 4E) and reduced expression of the mature neuron marker Calbindin (Fig. 4F). These data support the notion that increasing age is associated with a delay in maturation of adult-born neurons (Trinchero et al., 2017).

Intriguingly, recombined neurons in 24-week old β-cat^ex3^ iDCX mice featured a more mature dendritic arbor with longer dendrites and higher number of branch points compared to control neurons already three days after recombination (Fig. 5B and C). Thirteen days after-recombination, dendrite morphology was comparable between experimental groups (Fig. 5C).

**Fig. 5.**
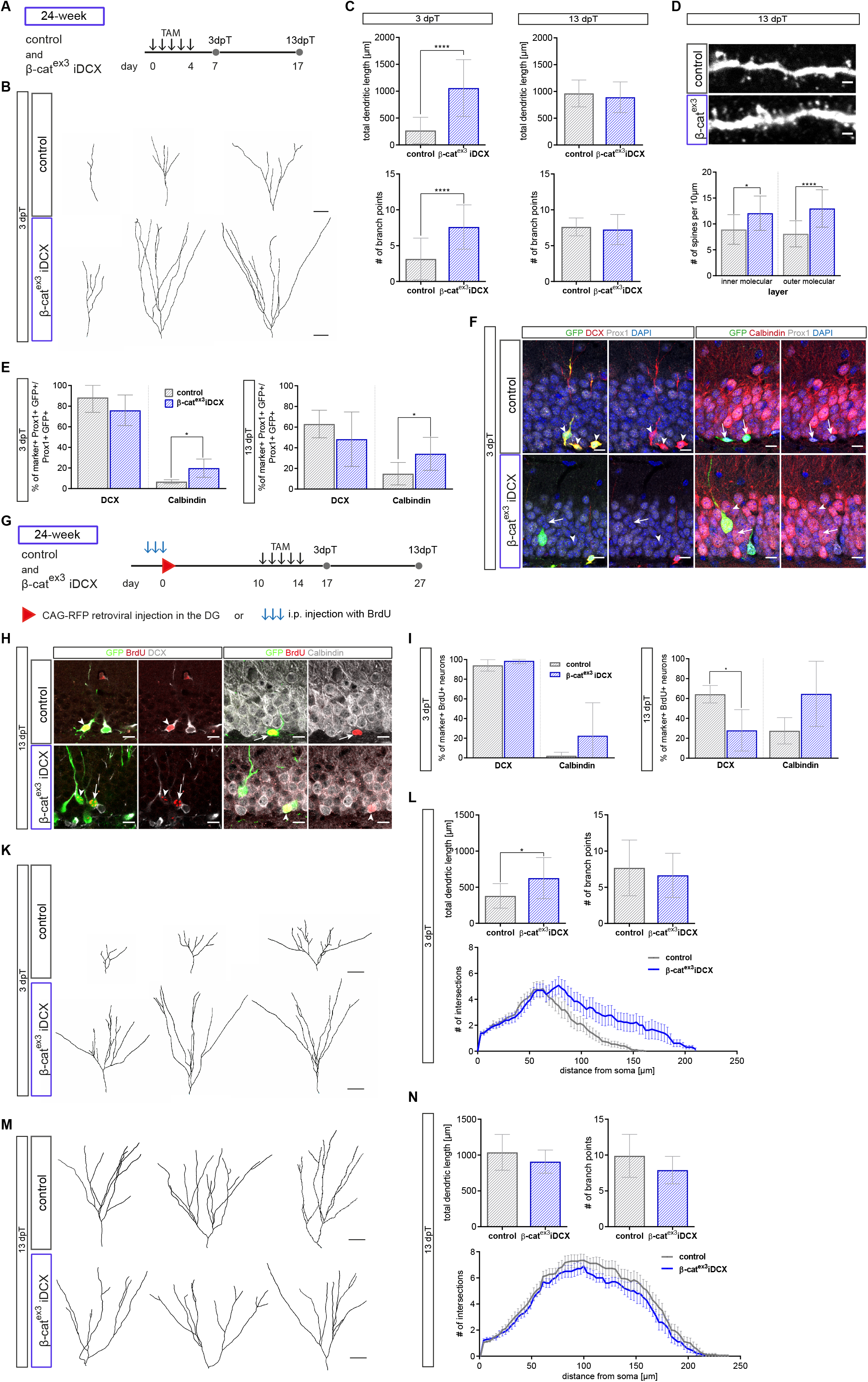
Precocious activation of canonical Wnt-signaling in middle-aged mice rescues aging phenotype. **A** Experimental scheme to analyze the impact of canonical Wnt-signaling on adult-born neuron development in middle-aged mice. Tamoxifen (TAM) was applied i.p. every 12 hours for five days. Animals were analyzed 3 and 13 days post Tamoxifen (dpT). **B** Representative reconstructions of control and β-cat^ex3^ iDCX neurons in 24-week old mice at 3dpT. Scale bar= 10μm **C** Dendritic length and number of branch points was increased in 24-week-old β-cat^ex3^ iDCX compared to 24-week-old controls at 3 dpT (control: n= 21 cells from 5 animals, β-cat^ex3^ iDCX: n= 20 cells from 6 animals). Dendritic length and number of branch points were comparable at the 13 dpT time point (control: n= 21 cells from 5 animals, β-cat^ex3^ iDCX: n= 20 cells from 8 animals). **D** Representative images showing dendritic segment of the inner molecular layer of the DG in 24-week old mice at 13dpT. Quantification of the number of spines showed increased spine density in inner molecular and outer molecular layer in 24-week old β-cat^ex3^ iDCX mice at 13dpT (control n= 14 cells from 4 animals, β-cat^ex3^ iDCX: n= 20 cells from 8 animals). Scale bars= 1μm. **E** Quantification of marker expression. The fraction of Calbindin expressing cells is increased in the recombined population in β-cat^ex3^ iDCX at 3dpT and 13dpT (3dpT: control: n= 4 animals, β-cat^ex3^ iDCX: n= 9 animals, 13 dpT: control: n= 8 animals, β-cat^ex3^ iDCX: n= 10 animals). **F** Representative pictures of recombined (GFP+) neurons in 24-week old animals co-expressing Prox1 as a marker for neuronal fate and the stage specific markers DCX for immature neurons and Calbindin for mature neurons. Arrows and arrowheads indicate marker negative and marker positive cells, respectively. Scale bar= 10μm **G** Experimental scheme for birthdating of adult-born neurons prior to recombination in 24-week old mice. For morphology analysis, CAG-RFP was stereotactically injected into the DG (K-M); for marker expression analysis, BrdU was injected i.p. every 24 hours for three days (H-I). Tamoxifen was applied i.p. every 12 hours for five days. Mice were sacrificed 3dpT and 13dpT. **H** Representative pictures of recombined (GFP+) BrdU-labeled neurons in 24-week old control and β-cat^ex3^ iDCX animals expressing the stage specific markers DCX for immature neurons and Calbindin for mature neurons. Arrows and arrowheads indicate marker negative and marker positive cells, respectively. Scale bar= 10μm **I** Quantification of DCX and Calbindin expression in BrdU-labeled recombined cells. At 13dpT, labeled neurons in β-cat^ex3^ iDCX animals show a shift towards a more mature marker profile. (3dpT: control: n= 4 animals, β-cat^ex3^ iDCX: n= 8 animals 13dpT control: n= 5 animals, β-cat^ex3^ iDCX: n= 5 animals). **K** Representative reconstructions of RFP-birthdated control and β-cat^ex3^ iDCX neurons in 24-week old mice at 3dpT. Scale bar= 30μm **L** Quantification of dendritic length, branch points, and Sholl analysis showed increase in complexity for birthdated β-cat^ex3^ iDCX cells at 3dpT (control: n= 22 cells from 4 animals, β-cat^ex3^ iDCX: n= 14 cells from 5 animals). **M** Representative reconstructions of RFP-birthdated control and β-cat^ex3^ iDCX neurons in 24-week old mice at 3dpT. Scale bar= 30μm **N** Quantification of dendritic length, branch points, and Sholl analysis displayed no difference between β-cat^ex3^ iDCX and control cells at 13dpT (control: n= 18 cells from 4 animals, β-cat^ex3^ iDCX: n= 14 cells from 3 animals) Data represented as mean ± SD (Sholl analysis: mean ± SEM), significance was determined using two-way ANOVA for Sholl analysis and two-tailed Mann-Whitney U Test for all other analyses, significance levels were displayed in GP style [p<0.0332 (*), p<0.0021 (**) and p<0.0002 (***), p<0.0001 (****)].

Neurons in β-cat^ex3^ iDCX mice, however, featured dendritic spine densities that were higher than those of neurons in middle-aged control animals (Fig. 5D) and that were comparable to spine densities of neurons in young adult control mice (SI Appendix, Fig. S1A). In addition, increased β-catenin activity resulted in a shift towards the expression of a mature marker profile. Three days after recombination, the β-cat^ex3^ iDCX group showed a trend towards lower DCX levels (control: 88%; β-cat^ex3^ iDCX: 76%) and a small but significant increase in the fraction of recombined neurons (GFP+ Prox1+) expressing Calbindin (control: 7%; β-cat^ex3^ iDCX: 20%) (Fig. 5E and F). Thirteen days after recombination, this shift towards a more mature marker profile was even more pronounced with 48% of recombined neurons expressing DCX and 34% expressing Calbindin in β-cat^ex3^ iDCX animals (control: DCX+ 63%; Calbindin+ 15%) (Fig. 5E). Notably, the proportion of mature Calbindin+ neurons in middle-aged β-cat^ex3^ iDCX animals was highly similar to the proportion of Calbindin+ neurons in young adult mice (Fig. EV1B).

To further substantiate accelerated neuronal maturation in middle-aged β-cat^ex3^ iDCX animals, we compared the developmental trajectory of neurons of the same birthdate between experimental conditions. Birthdating with BrdU or retrovirus was performed at the age of 24 weeks and recombination was induced from day 10 until day 14 after the birthdating procedure. Recombined cells were identified on the basis of GFP expression. Three days after recombination, β-cat^ex3^ iDCX animals showed a trend towards a higher fraction of Calbindin+ neurons amongst BrdU-labeled neurons. Thirteen days after recombination the switch towards the expression of a mature marker profile became obvious: thus, β-cat^ex3^ iDCX animals showed a significant downregulation of DCX (β-cat^ex3^ iDCX: 28 %; control: 64 %) and a substantial upregulation of Calbindin (β-cat^ex3^ iDCX: 65 %; control: 28%) in recombined neurons (Fig. 5H and I). Notably, the fraction of Calbindin expressing neurons in 24-week old β-cat^ex3^ iDCX reached levels that were comparable to levels in young adult control mice (Fig. EV1C). Next, we analyzed the morphology of CAG-RFP retrovirus labeled neurons. Three days after recombination, neurons with activated canonical Wnt/β-catenin-signaling displayed a substantially more mature dendrite morphology with increased total dendrite length (Fig. 5K and L). Thirteen days after recombination, neurons in both experimental groups featured a dendritic arbor that was highly comparable to the dendritic arbor of neurons of the same birthdate generated in young adult mice (Fig. 5M and N; Fig. EV1D). Collectively, these data demonstrate that increasing canonical Wnt/β-catenin-signaling activity counters the age-associated protracted maturation of adult-born dentate granule neurons.

## Discussion

Our study identifies activity of canonical Wnt/β-catenin-signaling as a cell-autonomous that timing and dosage of β-catenin-signaling regulate the tempo of dendritogenesis and the establishment of the highly stereotypic dendritic arbor of dentate granule neurons. Moreover, our data uncovers impaired canonical Wnt-signaling activity as a target to alleviate the age-associated maturation defect of adult-born neurons.

Wnt-dependent signaling pathways serve pleiotropic functions in embryonic and adult neurogenesis. While canonical Wnt-signaling has been primarily linked to early neurodevelopmental processes such as precursor proliferation and fate determination (Chenn & Walsh, 2002, Hirabayashi, Itoh et al., 2004, Kuwabara et al., 2009, Lie et al., 2005, Woodhead, Mutch et al., 2006), non-canonical Wnt-signaling pathways are considered the main drivers of neural circuit formation and plasticity (He, Liao et al., 2018, McLeod & Salinas, 2018, Salinas, 2012). We demonstrate that dendritogenesis of adult-born dentate granule neurons is dependent on canonical Wnt/β-catenin-signaling, which complements recent evidence that canonical Wnt/β-catenin-signaling contributes to neuronal maturation and circuit formation in embryonic neurodevelopment (Martin et al., 2018, Ramos-Fernandez, Tapia-Rojas et al., 2019, Viale, Song et al., 2019).

We found that in adult neurogenesis, genetic inhibition and age-associated decrease of Wnt/β-catenin-signaling activity were accompanied by a morphologically immature dendritic arbor and delayed dendritic development, respectively; whereas enhanced β-catenin activity by induction of the β-cat^ex3^ transgene countered the age-associated delay in dendrite development, mature neuronal marker expression and spine formation in middle-aged mice. These data strongly suggest that activity of the Wnt/β-catenin-signaling pathway modulates the tempo of dendrite development and neuronal maturation and that the age-associated retardation of adult-born neuron maturation is caused by dampened Wnt/β-catenin-signaling activity.

We cannot exclude that the β-cat^ex3^ driven rescue of adult-born neuron maturation in middle-aged mice was at least in part caused by non-transcriptional functions of β-catenin (Yu & Malenka, 2003). The observation that inhibition of β-catenin-signaling driven transcriptional activity via dnLEF resulted in an immature neuronal morphology together with the finding that canonical Wnt-signaling reporter activity increases during adult-born neuron maturation, however, argues that β-catenin driven transcription has a significant impact on dendritic growth. How Wnt/β-catenin-signaling regulates dendritogenesis and spine formation remains to be determined. Neurotrophin 3 was recently suggested to mediate dendritic growth and synapse formation downstream of β-catenin-signaling during embryonic neurogenesis (Viale et al., 2019). Given the ability of neurotrophins to modulate dendritogenesis in adult hippocampal neurogenesis (Bergami et al., 2008, Chan, Cordeira et al., 2008, Trinchero et al., 2017), it is tempting to speculate that a similar Wnt/β-catenin-signaling / neurotrophin axis also operates during dendritogenesis of adult-born neurons.

In contrast to middle-aged mice, in which enhanced β-catenin activity rejuvenated the developmental trajectory and produced neurons with a normal dendritic morphology and physiological spine densities, increased β-catenin activity in young mice transiently accelerated dendritic growth but ultimately resulted in a stunted dendritic arbor and increased spine densities. We speculate that the differences in outcome between young adult and middle-aged mice are caused by age-associated differences in the tone of Wnt/β-catenin-signaling in the neurogenic lineage. Middle-aged mice have reduced Wnt/β-catenin-signaling in maturing neurons and induction of the β-cat^ex3^ transgene may restore pathway activity to levels found in young adult mice. In contrast expression of the β-cat^ex3^ transgene in maturing neurons of young adult mice may result in excessive Wnt/β-catenin-signaling activity, which disrupts physiological dendritogenesis and spine development, which would be in line with the observation that restriction of Wnt/β-catenin-activity via a primary cilia-dependent mechanism is essential for dendritic refinement and correct synaptic integration of adult-born neurons (Kumamoto, Gu et al., 2012).

A key finding of this study is the demonstration that an ON-OFF-ON pattern of Wnt/β-catenin-signaling activity parallels neurogenesis in the adult dentate gyrus and that not only the re-activation of canonical Wnt-signaling but also its initial attenuation are essential for dendritogenesis. Why canonical Wnt-signaling activity has to be downregulated during early phases of neurogenesis remains to be determined. Given the evidence for crosstalk and functional antagonism between different Wnt-pathways (Baksh, Boland et al., 2007, Mentink, Rella et al., 2018), we speculate that sustained β-catenin-signaling interferes with non-canonical Wnt-signaling pathways and their essential function in dendrite patterning and growth (Arredondo et al., 2019, Goncalves et al., 2016a, Schafer et al., 2015).

The mechanism underlying reactivation of Wnt/β-catenin-signaling remains to be determined. Down-regulation of canonical signaling components was shown to drive the attenuation of Wnt/β-catenin-signaling activity in the early neurogenic lineage (Schafer et al., 2015), Increasing expression of canonical signaling components may allow the maturing neuron to regain responsiveness to canonical Wnt-ligands in the dentate gyrus (Gogolla, Galimberti et al., 2009, Lie et al., 2005). Another consideration is that the adult-born neuron encounters new environments during its development. While the cell body remains in the dentate gyrus and is continuously exposed to the same set of signals, the dendritic and axonal compartments gain access to potential new sources of Wnt-ligands during their growth, such as the molecular layer, the hilus and the CA3 region, which may trigger an increase in Wnt/β-catenin-signaling activity.

Aging, neurodegenerative and -psychiatric diseases impede on the functional integration of adult-born hippocampal neurons (Fitzsimons et al., 2013, Kim et al., 2012, Li et al., 2009, Llorens-Martin et al., 2015, Sun et al., 2009, Trinchero et al., 2017, Winner et al., 2011). Considering the powerful modulation of dendrite and spine development by Wnt/β-catenin-signaling activity the question arises whether aberrant Wnt/β-catenin-signaling activity contributes to these pathologies and can be targeted to improve neurogenesis-dependent hippocampal plasticity. Our finding that the age-associated protraction of adult-born neuron maturation is paralleled by a substantial drop in canonical Wnt-signaling activity and that activation of Wnt/β-catenin-signaling rejuvenated the time-line of maturation-associated marker expression, dendrite growth and spine formation in middle-aged mice, renders Wnt/β-catenin-signaling a promising candidate to ameliorate age-related impairment of hippocampal function.

## Material and Methods

### Experimental model and subject details

All experiments were carried out in accordance with the European Communities Council Directive (86/609/EEC) and were approved by the governments of Upper Bavaria and Middle-Franconia. DCX::CreER^T2^ mice (Zhang et al., 2010), Ctnnb1^(ex3)fl^ mice (Harada et al., 1999), BATGAL mice (Maretto et al., 2003), Axin2^LacZ^ mice (Lustig et al., 2002) and CAG-CAT-GFP mice (Nakamura et al., 2006) have been described previously. For all experiments, mice were grouped housed in standard cages with ad libitum access to food and water under a 12h light/dark cycle. DCX::CreER^T2^; CAG-CAT-GFP; Ctnnb1^(ex3)fl^ and DCX::CreERT2; CAG-CAT-GFP; Ctnnb1^(ex3)wt^ animals were initially generated from the same cross, and subsequently maintained as separate lines in the following to enable homozygous breeding. Male and female mice were used for experiments.

### Method details

#### Tissue processing

For brain tissue collection, mice were anesthetized using CO2 and transcardially perfused with phosphate-buffered saline (PBS, pH7.4) for five minutes at a rate of flow of 20 ml per minute followed by fixation with 4% paraformaldehyde (PFA) in 0.1 mM Phosphate Buffer (pH 7.4, Roth, Cat# 0335) for five minutes at a rate of 20 ml/min. Brains were post-fixed in 4% PFA at 4°C overnight and subsequently dehydrated in 30% sucrose solution. Frozen brains were either coronally or sagittally cut using a sliding microtome (Leica Microsystems, Wetzlar, Germany). Sections were stored at −20°C in 96-well plates, filled with 200 μl cryoprotection solution per well.

#### Genotyping

The following primers were used for genotyping DCX::CreER^T2^, CAG-CAT-GFP, Ctnnb1^(ex3)fl^ and BATGAL mice:

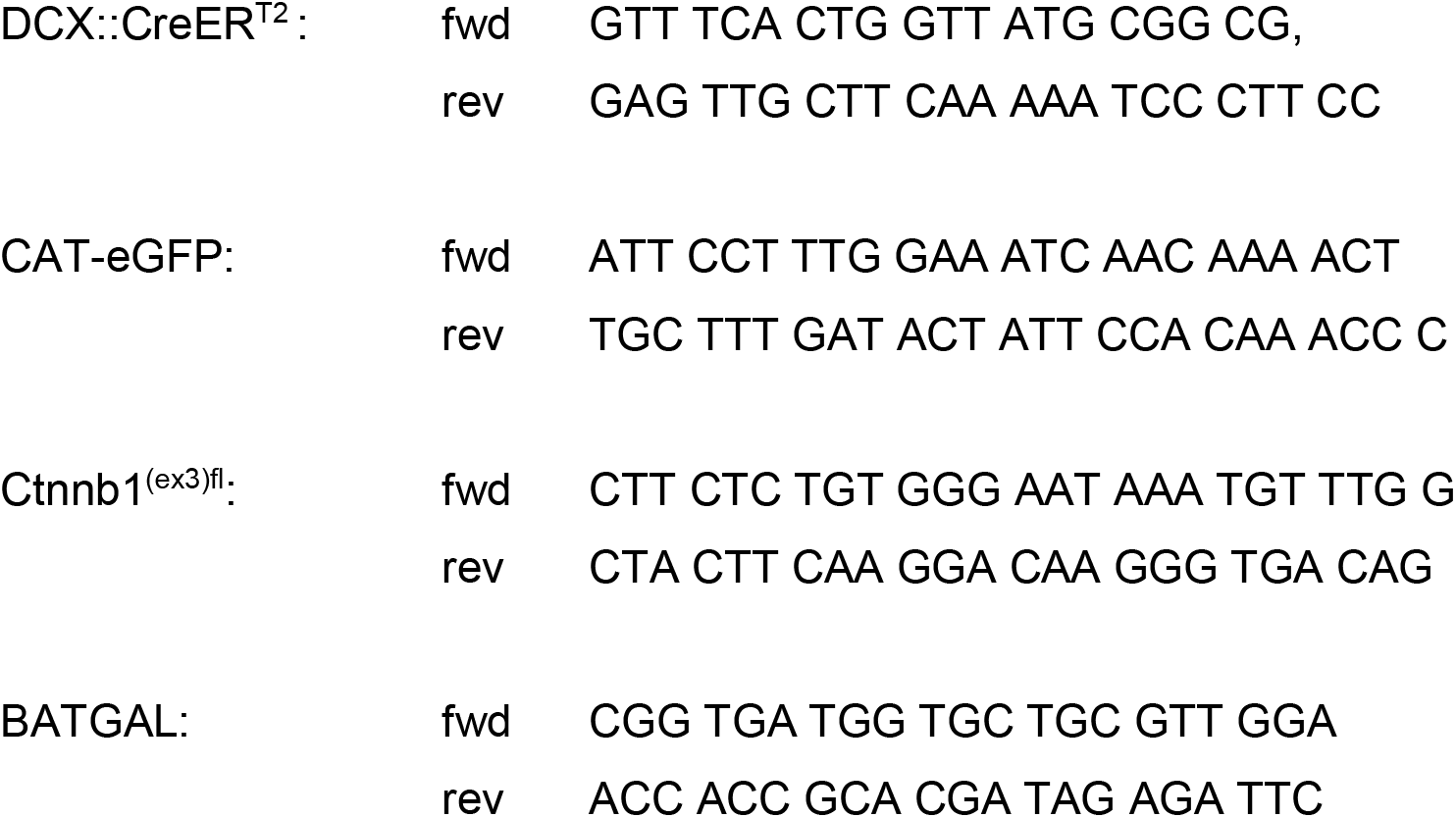

#### Tamoxifen administration

Tamoxifen (Sigma, Cat# T5648) was dissolved at a concentration of 10 mg/ml in ethanol and sunflower seed oil under constant agitation at room temperature. To induce recombination, 8 week-old and 24 week-old animals were intraperitoneally (i.p.) injected with 1 mg Tamoxifen every twelve hours for five consecutive days (Mori, Tanaka et al., 2006).

#### BrdU administration

Bromodeoxyuridine (BrdU, Sigma-Aldrich, Cat# B5002) was dissolved in sterile 0.9% NaCl solution at a concentration of 10 mg/ml. For birthdating experiments 8-week old, 24-week old and 36-week old animals were intraperitoneally injected with 100 mg/kg BrdU three times: i) every two hours for the 30-minute time point [described as 0 days post injection (dpi) in figures] ii) every 24 hours for all other time points.

#### Histology and counting procedures

Immunofluorescence stainings were performed on 40μm and 80μm thick free-floating brain slices. Selected brain slices were washed five times for ten minutes with Tris-buffered saline (TBS) in netwell inserts at room temperature (RT). Blocking and permeabilization was conducted in blocking solution (3% donkey serum and 0.25% Triton X-100 in TBS) for one hour at RT followed by incubation with the primary antibodies in blocking solution at 4°C for 72h.

After rinsing five times in TBS for 10 minutes at RT, brain slices were incubated with secondary antibodies in blocking solution at 4°C overnight. After rinsing three times for 10 minutes in TBS at RT sections were incubated with 4?,6-Diamidin-2-phenylindol (Dapi, Sigma, Cat# D9542) for 10 minutes at RT and washed once for 10 minutes in TBS. Brain slices were mounted on slides and coverslipped with Aqua Poly/Mount (Polysciences, Cat# 18606). Object slides were stored at 4°C in the dark.

If BrdU-incorporation into the DNA was to be examined, additional pretreatment in 2N HCl was performed for 10 minutes at 37°C after staining for the other used antibodies was completed and fixated with 4%PFA for 10 minutes at RT. The slices were then incubated in 0.1M Borate buffer for 10 minutes at RT and rinsed three times in TBS. Fluorescent staining for BrdU antibody was then performed as described above.

Primary antibodies were visualized with Alexa-conjugated secondary antibodies (all 1:1000; Invitrogen). To amplify the GFP reporter signal biotinylated secondary antibody (1:500; Vector Laboratories) was incubated at 4°C overnight. After rinsing the sections 5 times for 10 minutes with TBS at RT, Fluorophore-conjugated streptavidin (Invitrogen) in blocking solution was incubated overnight at 4°C and the staining procedure was finished as described above.

Confocal single plane images and z-stacks were taken with a Zeiss LSM 780 confocal microscope (Carl Zeiss, Oberkochen, Germany) equipped with four laser lines (405, 488, 559 and 633nm) and 25x, 40x, and 63x objective lenses. As a standard, the number of total pixels per image and color depth was set to 1024 × 1024 and 16bit respectively. For co-expression analysis, the 25x oil immersion objective was used; morphology was analyzed using the 40x and 63x oil immersion objective. Z-stack step size for co-expression and morphology analyses was set to 1.5μm and 0.3μm, respectively. Images were processed using Fiji ImageJ. 3D reconstructions were obtained using Imaris software (Bitplane AG, Zürich, Switzerland).

#### Primary antibodies

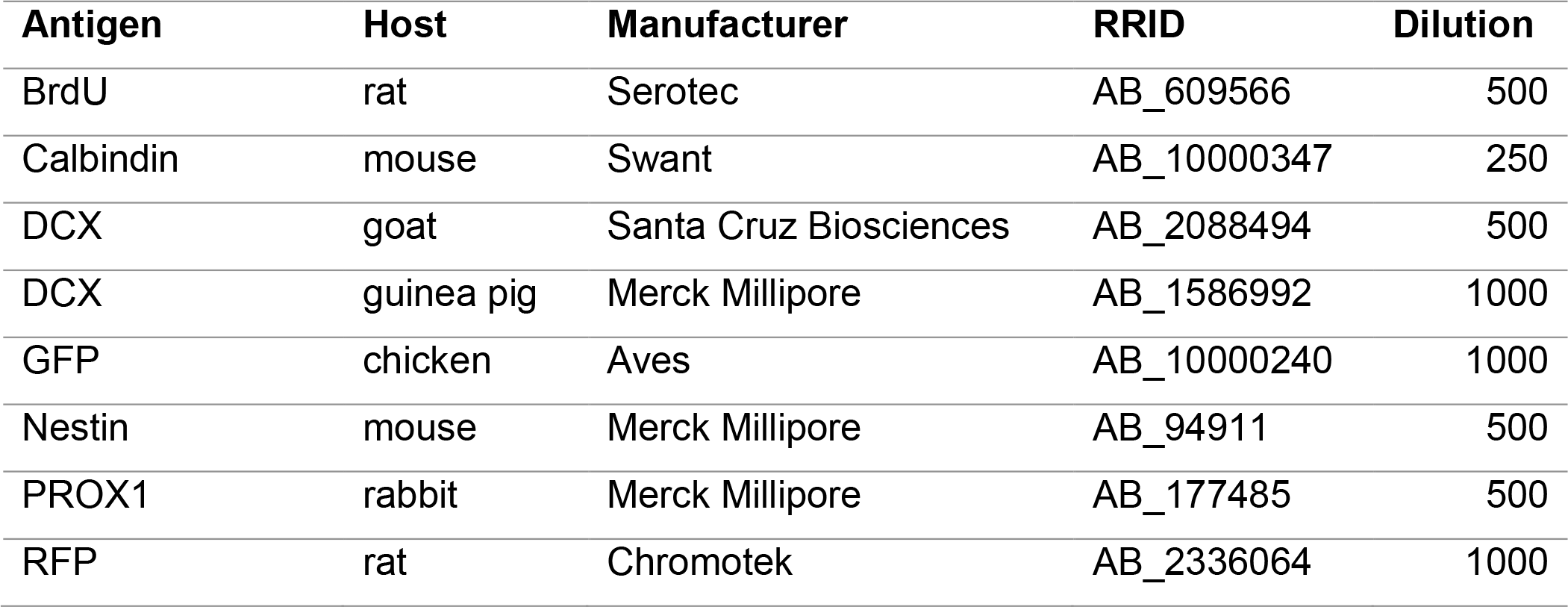
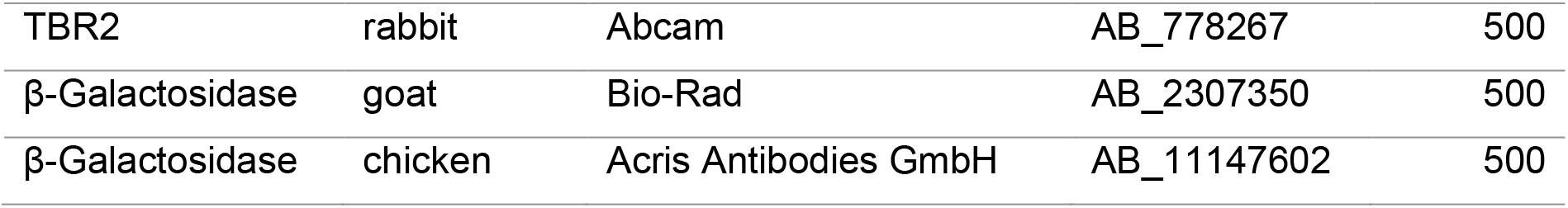

#### Retrovirus preparation and stereotactic injections

The retroviral plasmids CAG-GFP-IRES-Cre, CAG-dnLEF-IRES-GFP and the CAG-RFP have been described (Karalay et al., 2011, Steib, Schaffner et al., 2014, Tashiro, Sandler et al., 2006). Production of replication incompetent MML-retroviruses was performed in human 293-derived retroviral packaging cell line (293GPG) 1F8 cells as described previously (Tashiro, Zhao et al., 2006).

For stereotactic injections, 8-week-old and 24-week-old mice under running conditions were deeply anesthetized by injecting 300μl of a mixture of Fentanyl (0.05 mg/kg; Janssen-Cilag AG, New Brunswick, USA), Midazolam (5 mg/kg; Dormicum, Hoffmann-La Roche, Basel, Switzerland) and Medetomidine (0.5 mg/kg; Domitor, Pfizer Inc., New York, USA) dissolved in 0.9% NaCl. Mice were fixed in a stereotactic chamber and the scalp was opened. Holes were drilled into the skull for injections into both hemispheres (coordinates from bregma were −1.9 anterior/posterior, ±1.6 medial/lateral, −1.9 dorsal/ventral from dura). 900nl virus particle suspension diluted with PBS to a concentration of 2×10^8^ colony forming units per μl was injected into the DG of each hemisphere at a speed of 250 nl/min using a Digital Lab Standard Stereotaxic Instrument. Anesthesia was antagonized after surgery by injecting a mixture of Buprenorphine (0.1 mg/kg, Temgesic, Essex Pharma GmbH, Munich, Germany), Atipamezol (2.5 mg/kg, Antisedan, Pfizer Inc., New York, USA) and Flumazenil (0.5 mg/kg; Anexate, Hexal AG, Holzkirchen, Germany) dissolved in 0.9% NaCl.

### Quantification and statistical analysis

#### Expression analysis of stage-specific markers

Coexpression analysis were conducted using ImageJ software. For cell stage marker (Nestin, Tbr2, DCX, Prox1, Calbindin) co-expression analysis, >100 cells per animal were analyzed from at least four different animals. Expression of BrdU in BATGAL and Axin2^LacZ/+^ mice was determined in at least two sections containing the hippocampus of at least four different animals and for control and β-cat^ex3^ iDCX mice one section containing the hippocampus of at least five different animals. The number of biological replicates (n) analyzed is specified in the figure legends.

#### Dendritic and spine morphology analyses

To analyze detailed cell morphology, confocal images were taken using Zeiss LSM 780 confocal microscope (Carl Zeiss, Oberkochen, Germany) with a 40x and 63x oil immersion objective. For spine analysis the digital zoom was set to 3 enabling better spatial resolution. Brain slices for dendritic morphology analysis and spine analysis were 80-100 μm and 40 μm thick respectively. 3D reconstructions of neurons were obtained with Imaris using the Filament Tracer tool and the Surface Tracer tool. Values for total dendritic length, number of Sholl intersections, number of branch points and basal dendrites were exported. The number of spines were investigated using Fiji.

#### Statistical analysis

GraphPad prism was used for statistical analysis. The statistical significance level α was set to 0.05. Gaussian distribution was tested using the D’Agostino-Pearson omnibus test. If not applicable non-Gaussian distribution was assumed. Statistical significance was determined using the two-tailed Mann-Whitney U Test or two-way ANOVA for Sholl analysis and significance levels were displayed in GP style [p<0.0332 (*), p<0.0021 (**) and p<0.0002 (***), p<0.0001 (****)]. Unless otherwise stated in the figure legend, results are represented as mean ± SD and as mean ± SEM for Sholl analysis. The number of biological replicates (n) is specified for each analysis in the figure legend. For marker expression experiments n equals the number of individual animals analyzed and for morphology analysis n equals the number of neurons analyzed from a minimum of three different animals.

## Acknowledgments

We thank S. Jessberger and all members of the Lie laboratory for helpful discussions and comments on the manuscript. This work was supported by grants from the German Research Foundation (LI 858/6-3 and 9-1 to D.C.L, INST 410/45-1 FUGG), the Bavarian Research Network “ForIPS” and “ForINTER” to D.C.L., the University Hospital Erlangen (IZKF grants E12, E16, E21 to D.C.L.). J.H. and M.T.W. are members of the research training group 2162 “Neurodevelopment and Vulnerability of the Central Nervous System” funded by the Deutsche Forschungsgemeinschaft (270949263/ DFG GRK2162/1). The authors declare no competing financial interests.

## Author contributions

Conceptualization, J.H., N.P., D.C.L.; Investigation, J.H., M.-T.W., J.Z.; Formal analysis, J.H., M.-T.W., J.Z., D.C.L.; Resources and Funding acquisition, W.W., D.C.L.; Reagents, M.M.T., W.W.; Writing-Original draft, J.H., D.C.L.; Writing-Review and Editing, J.H., D.C.L.; Supervision: N.P., D.C.L.

## Expanded view figure legend

**Fig. EV1.**
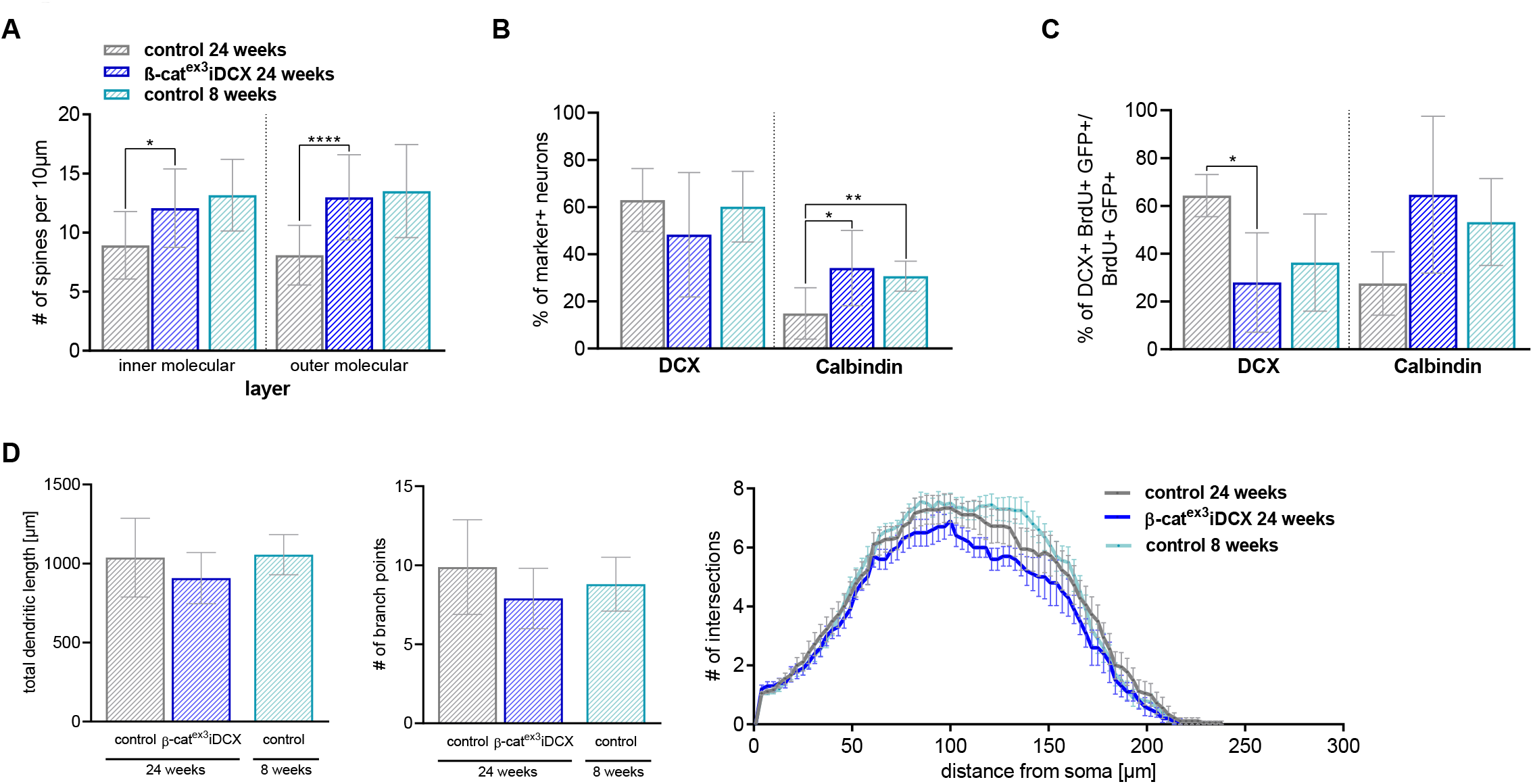
Induction of Wnt/β-catenin-signaling activity in middle-aged mice rejuvenates dendrite and spine development. **A** Quantative analysis of spine density at the inner molecular and outer molecular layer of the DG at 13dpT. The number of spines in 24-week-old β-cat^ex3^iDCX resembled the number of spines in 8-week-old control mice, indicating a rescue of delayed spinogenesis in middle-aged animals. **B** The fraction of neurons with DCX and Calbindin expression in middle-old β-cat^ex3^iDCX animals was comparable to the fraction in young control animals. **C** Birthdated neurons in middle-aged β-cat^ex3^iDCX animals and young control animals showed a comparable fraction of neurons expressing DCX and Calbindin at 13dpT. **D** 24-week-old control, β-cat^ex3^iDCX and young control animals have comparable dendritic parameters (total dendritic length, number of branch points, and Sholl analysis) at 13dpT. Data represented as mean ± SD(Sholl analysis: mean ± SEM), significance was determined using two-way ANOVA for Sholl analysis and two-tailed Mann-Whitney U Test for all other analyses, significance levels were displayed in GP style [p<0.0332 (*), p<0.0021 (**) and p<0.0002 (***), p<0.0001 (****)]

